# Conserving steppe-land birds under climate change: a gap analysis for the Eurasian Stone-curlew (*Burhinus oedicnemus*) in the Western Palearctic

**DOI:** 10.1101/2024.05.20.594951

**Authors:** Andrea Simoncini, Samuele Ramellini, Mattia Falaschi, Mattia Brambilla, Alexis Martineau, Alessandro Massolo, Dimitri Giunchi

## Abstract

Climate change is having dramatic impacts on the distribution of animals. Birds, and especially steppe-land birds, are particularly sensitive to climate change and identifying areas that are critical for their conservation is pivotal, as well as estimating the expected impact on these areas under different climate and land use change scenarios. In-situ climate refugia (areas suitable under both current and future climates) are especially valuable for the conservation of climate-sensitive species, and is therefore important to identify them and evaluate their coverage by protected areas. Via species distribution modelling, we aimed to identify in-situ climate refugia in the Western Palearctic for the Eurasian Stone-curlew *Burhinus oedicnemus*, an umbrella steppic species of conservation concern. We used a comprehensive dataset of occurrences in the breeding period to fine-tune a Maxent species distribution model and project it under three carbon emission scenarios of increasing severity for the year 2050. We then identified in-situ climate refugia and performed a gap analysis estimating the percentage of refugia falling within the network of currently protected areas. In all modelled future scenarios a northward expansion of suitable breeding habitats was predicted, and suitable areas had similar extents, with a slight increase of the overall suitability under more severe scenarios. According to our results, the Eurasian Stone-curlew has the potential to maintain viable populations in the Western Palearctic, even though dispersal limitations might hinder the colonization of newly suitable breeding areas. In-situ climate refugia were mainly identified outside protected areas, particularly in Northern Africa and the Middle East. Therefore, we advocate targeted actions in climate refugia to promote the conservation of this and other steppe-land species under global environmental change.

## 1. INTRODUCTION

Anthropogenic global change exerts a strong influence on organisms’ distribution, life-history, population viability, and thus, on their likelihood of extinction (Chen et al., 2011; Parmesan, 2006; Román-Palacios and Wiens, 2020; Selwood et al., 2015). One example is the shift in the distribution range following climate change (VanDerWal et al., 2013). Such distributional shifts have been described both for sessile and highly mobile animals (Williams and Blois, 2018). Birds, in particular, are excellent model organisms to understand the response of organisms to climate change, due to the presence of long-term datasets covering wide spatial scales (Brlík et al., 2021). Furthermore, in birds it has been shown that climate change can affect habitat suitability (Barbet-Massin et al., 2012a), which in turn affects population trends (Green et al., 2008). Steppe-land species are particularly sensitive to global change. In fact, such species experienced particularly dramatic declines, which were primarily linked to both agricultural intensification and afforestation following land abandonment (Burfield, 2005; Onrubia and Andrés, 2005; Silva et al., 2024). Previous studies using species distribution models (SDMs, Guisan and Zimmermann, 2000) to derive habitat suitability for steppe-land birds reported poleward range shifts and the persistence of large suitable areas according to global change scenarios (Estrada et al., 2016; Kiss et al., 2020). The Eurasian Stone-curlew *Burhinus oedicnemus* (Linnaeus, 1758; hereafter Stone-curlew) is a wide-ranging steppe-land species occurring in (pseudo-)steppes and farmlands in the Palearctic (Vaughan and Vaughan Jennings, 2005). This species is considered both an umbrella and flagship species (Caro, 2010; Hunter Jr et al., 2016), and its conservation can therefore benefit steppe-land species and, more generally, steppe-land ecosystems (Hawkes et al., 2019). The Stone-curlew suffered a severe population decline (>30%) in the second half of the 20^th^ century, mainly due to agricultural intensification (BirdLife International, 2018), but in recent decades positive trends have been reported in some areas of its range such as France (BirdLife International, 2017). It is now classified as Least Concern by the IUCN (BirdLife International, 2021), but information on the status of its populations is limited, and positive trends might be biased by the increased monitoring effort (Gaget et al., 2019). Indeed, the species is still considered of European conservation concern (SPEC3) and ongoing declines are reported for many regions of its range (BirdLife International, 2017; Gaget et al., 2019). Huntley et al. (2007) used SDMs to forecast the future distribution of the Stone-curlew under climate change scenarios, highlighting a northward shift of suitable areas. Still, their European-level analysis was limited to a coarse scale (∽ 50 km), hence preventing fine-scale applications such as comparisons with the current distribution of protected areas. To establish effective conservation practices, state-of-the-art habitat suitability scenarios for the focal species are required (McShea, 2014; Miller-Rushing et al., 2010). Rigorous analyses of prospective suitability dynamics are essential to reveal a species’ sensitivity to climate change and to guide conservation efforts (Stiels et al., 2021). In the spatial planning of the conservation of a species it is also important to define the areas currently suitable and predicted to remain so under all future climate scenarios, so-called ‘in-situ climate refugia’ (Brambilla et al., 2022). The identification of refugia, combined with a gap analysis, allows to detect areas critical for biological conservation that have been overlooked and/or ineffectively managed and is pivotal for planning conservation actions (Maiorano et al., 2006). Therefore, we aimed to: 1) describe the current availability of suitable breeding habitats, 2) provide a range of future breeding habitat suitability scenarios considering climatic changes, 3) identify in-situ climate refugia, and 4) assess the proportion of refugia within the current network of protected areas (gap analysis).

## 2. METHODS

### 2.1. Stone-curlew occurrences

We obtained a dataset of Stone-curlew occurrences during the breeding season (May-July; Vaughan and Vaughan Jennings, 2005) spanning twenty years (1996-2015). We retrieved data for the Western Palaearctic (*sensu* Snow et al., 1998) and excluded the two insular, genetically-distinct Macaronesian subspecies (*B. o. distinctus*, *B. o. insularum*). We downloaded data from eBird, Global Biodiversity Information Facility (gbif.org; https://doi.org/10.15468/dl.zxc278), ornitho.it, ornitho.cat and xeno-canto.org, as these proved useful in previous SDM studies (Avalos and Hernández, 2015; Coxen et al., 2017; Engelhardt et al., 2020; Ramellini et al., 2019). We also included survey data from the British Trust for Ornithology (in agreement with BTO’s data policy), data for Greece from the Ornithotopos database, the second European Breeding Bird Atlas (hereafter EBBA) and 2 × 2 km surveys in Greece (in formal agreement with the Hellenic Ornithological Society), and data for the Deux-Sèvres department from the Nature79 database (provided by Alexis Martineau). Finally, we considered nest points from Northern Italy collected by Dimitri Giunchi and collaborators in the period 2012-2015. We discarded observations outside the species’ current range (BirdLife International, 2022; Hagemeijer and Blair, 1997; Keller et al., 2020) to exclude vagrant individuals in non-breeding areas (Figure 1). We used point occurrences with a maximum spatial uncertainty of 5.5 km, which roughly approximates the species’ breeding home range size (Caccamo et al., 2011; Hawkes et al., 2021). This value (5.5 km) was also used to define the linear size of grid cells for all models developed in this study. Indeed, a model grain approaching home range or territory size is particularly desirable to obtain ecologically more meaningful models (Brambilla et al., 2024). Moreover, to reduce the spatial autocorrelation among occurrences and limit the effect of possibly biased sampling effort across space, we subsampled the dataset considering a minimum distance between occurrence points of 16 km, so that there were no occurrence points in adjacent cells. The final dataset consisted of 816 presences spread across the species’ range in the Western Palearctic (Figure 1).

**Figure 1.**
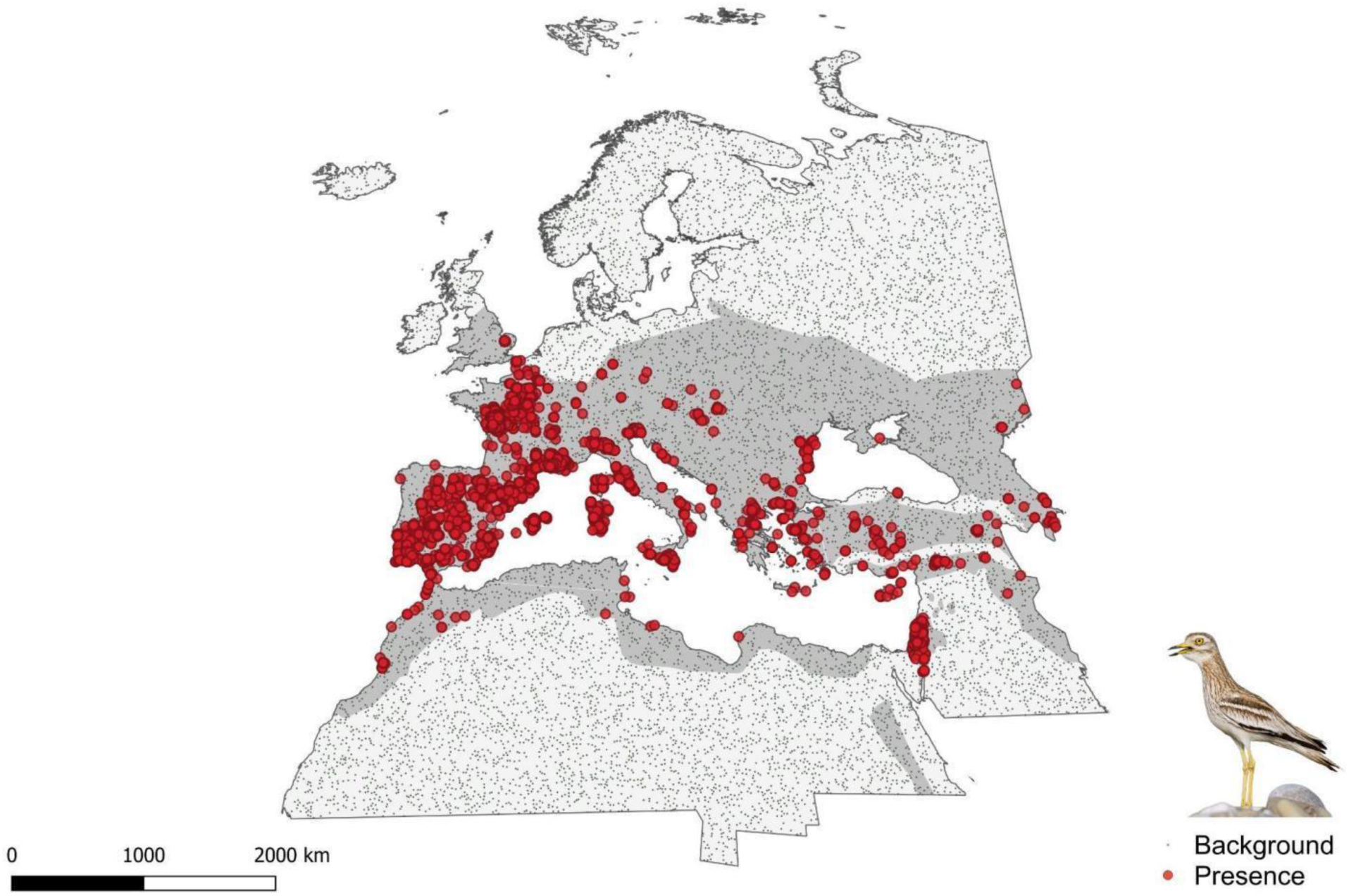
Data used to model Eurasian Stone-curlew (*Burhinus oedicnemus*) distribution. Occurrence and background points used to model breeding habitat suitability for the Eurasian Stone-curlew (*Burhinus oedicnemus*) in the Western Palearctic. Occurrences (red dots) and 10000 randomly selected background points (small dark dots) are shown. Data for the UK provided by the British Trust for Ornithology are not shown due to sharing restrictions to protect nesting sites. The projection used is World Mollweide (ESRI: 54009). The species’ range in the Western Palearctic, excluding the Macaronesian archipelago, is shown in dark grey and was obtained from the BirdLife International website (https://www.datazone.birdlife.org). Inset photo by Saverio Gatto.

### 2.2. Variable selection and preparation

We used an expert-based approach (Santini et al., 2021) to derive eleven biologically meaningful environmental predictors representing climate, land use and land cover (LULC). Climate is a major determinant of Stone-curlew’s distribution at the broad scale (Vaughan and Vaughan Jennings, 2005), as the species selects warm and dry areas to reproduce (Green et al., 2000; Keller et al., 2020; Vaughan and Vaughan Jennings, 2005). We therefore included three variables describing temperature (annual mean temperature, temperature of the warmest quarter, temperature seasonality) and three describing precipitation (annual precipitation, precipitation of the warmest quarter, precipitation seasonality). Climatic variables were obtained from CHELSAcruts (Karger and Zimmermann, 2018) at one year resolution. As ensuring a temporal match between predictors and occurrences increases the reliability of species-environment relationship (Gavrutenko et al., 2021), we computed the averages of climatic variables over the twenty-years period covered by observations (1996-2015). Climate stability can significantly affect the distribution of species, being one of the major drivers of diversification processes and endemism patterns (Rangel et al., 2018). We accounted for climate stability using the CSI (Climate Stability Index) index (Herrando-Moraira et al., 2022). LULC change has been shown to affect the distribution and abundance of the Stone-curlew (Burfield, 2005; Onrubia and Andrés, 2005). Stone-curlews exploit agricultural areas with low vegetation density for breeding and foraging (Caccamo et al., 2011; Vaughan and Vaughan Jennings, 2005). Arid and steppic grasslands are among the elective habitats for the species (Green et al., 2000; Hume and Kirwan, 2013; Teyar et al., 2021; Vaughan and Vaughan Jennings, 2005) as well as low shrubs, often mixed with grass and bare ground (Traba et al., 2013; Vaughan and Vaughan Jennings, 2005). Finally, breeding attempts in urbanized areas are increasingly reported (Biondi et al., 2015; Cutini et al., 2007; Giovacchini et al., 2017). We hence derived the percentage cover for the following classes: 1) agricultural, 2) grass, 3) shrub, and 4) urban. These predictors adequately represent the main LULC drivers of habitat suitability for the species previously described (Giovacchini et al., 2017; Vaughan and Vaughan Jennings, 2005). LULC variables were computed as the percentage cover of each class on the total surface of a raster grid cell, using the global land cover developed by the European Space Agency (European Space Agency, 2019) and computing the mean over the same twenty-years period of species’ occurrences (1996-2015). We tested for correlation among these variables using the Pearson linear correlation coefficients (Supplementary Materials S1) and excluded four climatic variables (temperature of the warmest quarter, temperature seasonality, precipitation of the warmest quarter, precipitation seasonality) to obtain a set of seven uncorrelated predictors with Pearson’s coefficient always below 0.7.

### 2.3. Future scenarios

We retrieved future climatic layers for five Intergovernmental Panel on Climate Change General Circulation Models (GCMs) that gained the minimum score in interdependence (Sanderson et al., 2015): CESM1-CAM5, FIO-ESM, IPSL-CM5A-MR, MIROC5 and MPI-ESM-MR. We considered three carbon emission scenarios corresponding to the IPCC’s Representative Concentration Pathways 2.6, 4.5 and 8.5 (RCP 2.6, 4.5 and RCP 8.5). This suite of RCPs is often selected as it represents a full spectrum of scenarios, including an optimistic (2.6), a moderate (4.5) and a pessimistic (8.5) scenario (Waynat et al., 2009). We produced bioclimatic variables for every GCM × RCP combination from the climatic variables available in the CHELSA CMIP-5 timeseries (Karger et al., 2017). Future variables represented predicted mean conditions over a twenty-years period (2041-2060). The future CSI index (a single raster for the future) was obtained from Herrando-Moraira et al., 2022. LULC variables were kept at their 2016 values, due to the extremely low correspondence between the dataset used to train models and any dataset of future LULC available to our knowledge. We also note that incorporating fixed LULC variables in future projections of species distributions increases their reliability as LULC is expected to change more slowly compared to climate and therefore it is useful to constrain suitable areas to those with LULC values within the niche of the species (Stanton et al., 2012). All variables were computed or resampled at the 5.5 × 5.5 km grid resolution, under the equal-area projection World Mollweide (ESRI: 54009).

### 2.4. Species distribution modelling workflow

Considering the inherent spatial bias in the Stone-curlew point occurrences, we opted for a machine learning algorithm using a maximum entropy (Maxent) approach (Phillips et al., 2006). In fact, this is considered among the best suited and robust methods for modelling species distribution as it allows to fine-tune models, while contextually balancing power and complexity (Radosavljevic and Anderson, 2014; Warren and Seifert, 2011). Maxent requires background points to describe the suite of environmental conditions found in the study area and compare them with conditions at occurrence points (Merow et al., 2013). Therefore, we considered 10000 background points randomly generated over the study area (Barbet-Massin et al., 2012b). Tuning is essential to reduce overfitting and identify appropriate hyperparameters (Brambilla et al., 2022; Radosavljevic and Anderson, 2014). Therefore, we tuned the regularization multiplier (beta, a penalty of all parameters included in Maxent models that affects model complexity) comparing eight values (from one to eight at incremental steps of one), and features, testing two different combinations: linear and quadratic (LQ) or linear, quadratic, and hinge (LQH). Linear and quadratic features have often been proved to be the most biologically meaningful (Anderson and Gonzalez, 2011; Syfert et al., 2013), whereas hinge features allow to account for more complex species-environment relationships (Merow et al., 2013). To compare the performance of different models we used the following metrics: 1) Area Under the Curve (AUC) (Fielding and Bell, 1997) of the Receiver Operating Characteristic (ROC) plot on a test dataset (AUC_test_), a measure of the predictive ability of a model that is threshold-independent, 2) the difference between the training and test AUC (AUC_diff_) (Radosavljevic and Anderson, 2014), as a measure of overfitting (with higher values indicating model overfitting), and 3) Continuous Boyce Index (CBI) (Boyce et al., 2002), a measure that is independent of species prevalence. Spatial autocorrelation can strongly bias inference on the factors determining species’ distribution, often leading to overfitting and overrated predictive performance of machine-learning algorithms (Ploton et al., 2020; Roberts et al., 2017). Therefore, in all tests of model performance, we used a four-fold spatial cross validation (Muscarella et al., 2014). This procedure splits the occurrences and background points in four spatial blocks maximising spatial independence, and then uses three blocks for model training and the remaining one for testing, iterating four times. We estimated environmental variable contribution in the best-performing model computing permutation importance, i.e., randomly permuting the values of a variable while keeping the others at their mean values and evaluating the drop in AUC_test_, expressed as percentage (Thuiller et al., 2009). After tuning the models, the one showing the best trade-off among performance metrics was used to predict current and future (2041-2060) breeding habitat suitability for the Stone-curlew in the Western Palearctic. For the choice of the model representing the best trade-off, even if we checked all metrics, we gave priority to models with a low overfitting given the need to extrapolate to future conditions. When projected to new time periods, SDMs might encounter environmental conditions that are not found in the calibration area (Elith et al., 2010; Zurell et al., 2012), producing spurious predictions (Owens et al., 2013). This occurs when: 1) a variable is outside the range found when training (i.e., strict extrapolation), or 2) each variable is within the calibration range, but the combination of predictors is new (i.e., combinatorial extrapolation) (Elith et al., 2010; Zurell et al., 2012). We used the environmental overlap mask to identify the areas where strict and combinatorial extrapolation occur (Zurell et al., 2012).

### 2.5. Gap analysis

Areas predicted as suitable under both current and future conditions (i.e., *in situ* climate refugia) are fundamental to plan conservation actions (Estrada et al., 2016; Thuiller et al., 2019). We binarized each habitat suitability map considering as suitable the cells with a suitability above the maximum training sensitivity plus specificity (habitat suitability > 0.3031) of the cloglog model output as this binarization is considered robust and reliable (Liu, 2012). These maps were then used to identify cells predicted to remain suitable under future scenarios (i.e., in-situ climate refugia). Then, we computed the percentage of climate refugia found within the World Database on Protected Areas, www.protectedplanet.net (downloaded on April 10^th^, 2023). From this database we removed protected areas considered as “proposed”, retaining only those currently established.

### 2.6. R packages

All analyses were performed in the R environment (version 4.3.1; R Core Team 2023) with the packages: ‘usdm’ v. 1.1-18 (Naimi, 2013), ‘humboldt’ v. 1.0.0 (Brown and Carnaval, 2019), ‘dismo’ v. 1.3.14 (Hijmans et al., 2017), ‘raster’ v. 3.6.20 (Hijmans, 2018), ‘ecospat’ v. 4.0.0 (Di Cola et al., 2017), ‘ENMeval’ v. 0.2.1 (Muscarella et al., 2014), ‘rmaxent’ v. 0.8.5 (https://github.com/johnbaums/rmaxent), ‘mecofun’ v. 0.5.1 (https://gitup.uni-potsdam.de/macroecology/mecofun).

## 3. RESULTS

### 3.1. Species distribution modelling

Based on the tuning of regularization multipliers and features, we selected as best the model with linear and quadratic features and beta = 1, as it showed the lowest overfitting (AUC_diff_ =0.057 ± 0.088 SD) while retaining excellent AUC_test_ (0.872 ± 0.081 SD) and CBI (0.990 ± 0.008 SD) metrics (Table 1). The tuned model provided a reliable description of the ecological niche of the Stone-curlew (Figure 2) and the current habitat suitability of the Western Palearctic for this species (Figure 3). The variable with the highest permutation importance was annual mean temperature, followed by agricultural cover, annual precipitation and climate stability (Table 2). Response curves (Figure 2) showed a unimodal relationship between the probability of Stone-curlew occurrence and annual mean temperature (with a peak between 15 and 20 °C), annual precipitation (with a peak between 800 and 1200 mm), and a quadratic relationship with agricultural cover. Maps of predicted habitat suitability for the year 2050 showed an increase of the suitable areas towards north-east under all scenarios, particularly under more severe scenarios (Figure 3). Slight decreases of breeding habitat suitability were predicted under all RCPs at the southern range border, e.g., in Northern Africa (Figure 3). The agreement between GCMs for future predictions was consistently high except for a band across Central Europe were uncertainty was higher under all scenarios (Figure 3). Strict extrapolation was extremely low under current conditions, whereas combinatorial extrapolation was slightly higher, although still confined to a few areas (Figure S1). Strict extrapolation was low under future scenarios for all GCMs but higher compared to the current, and it increased under more severe scenarios (Figures S2-S7). This mainly concerned the southern border of the Western Palearctic (Figures S2-S7). Combinatorial extrapolation had a similar pattern although it also occurred more consistently in the Middle East and Northern Europe (Figures S7-S11).

**Figure 2.**
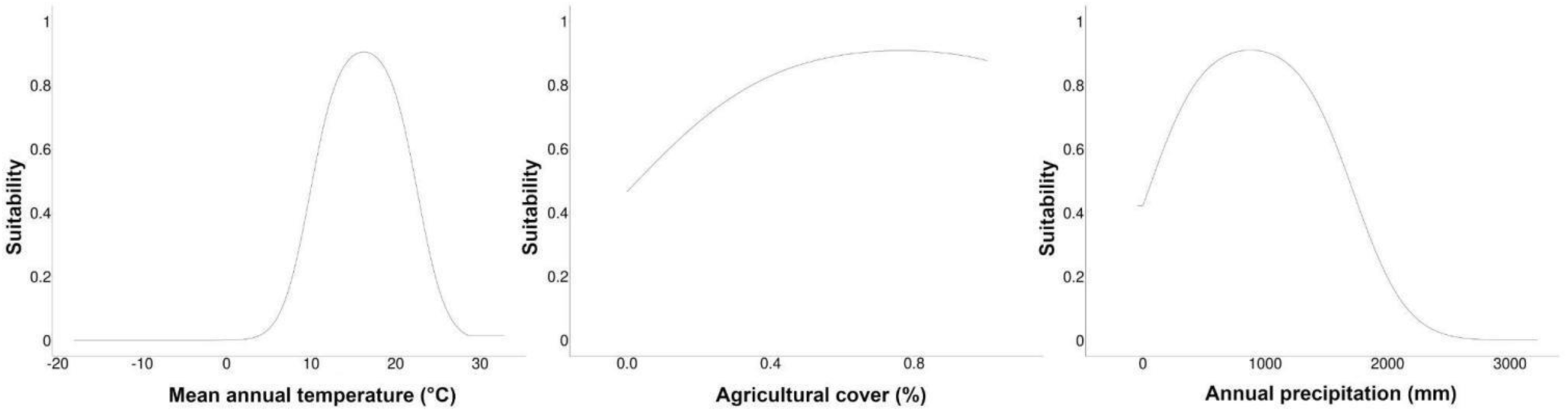
Response curves for a distribution model of Eurasian Stone-curlew (*Burhinus oedicnemus*). Response curves for the three variables with the highest permutation importance in a Maxent species distribution model for the Eurasian Stone-curlew (*Burhinus oedicnemus*) in the Western Palearctic.

**Figure 3.**
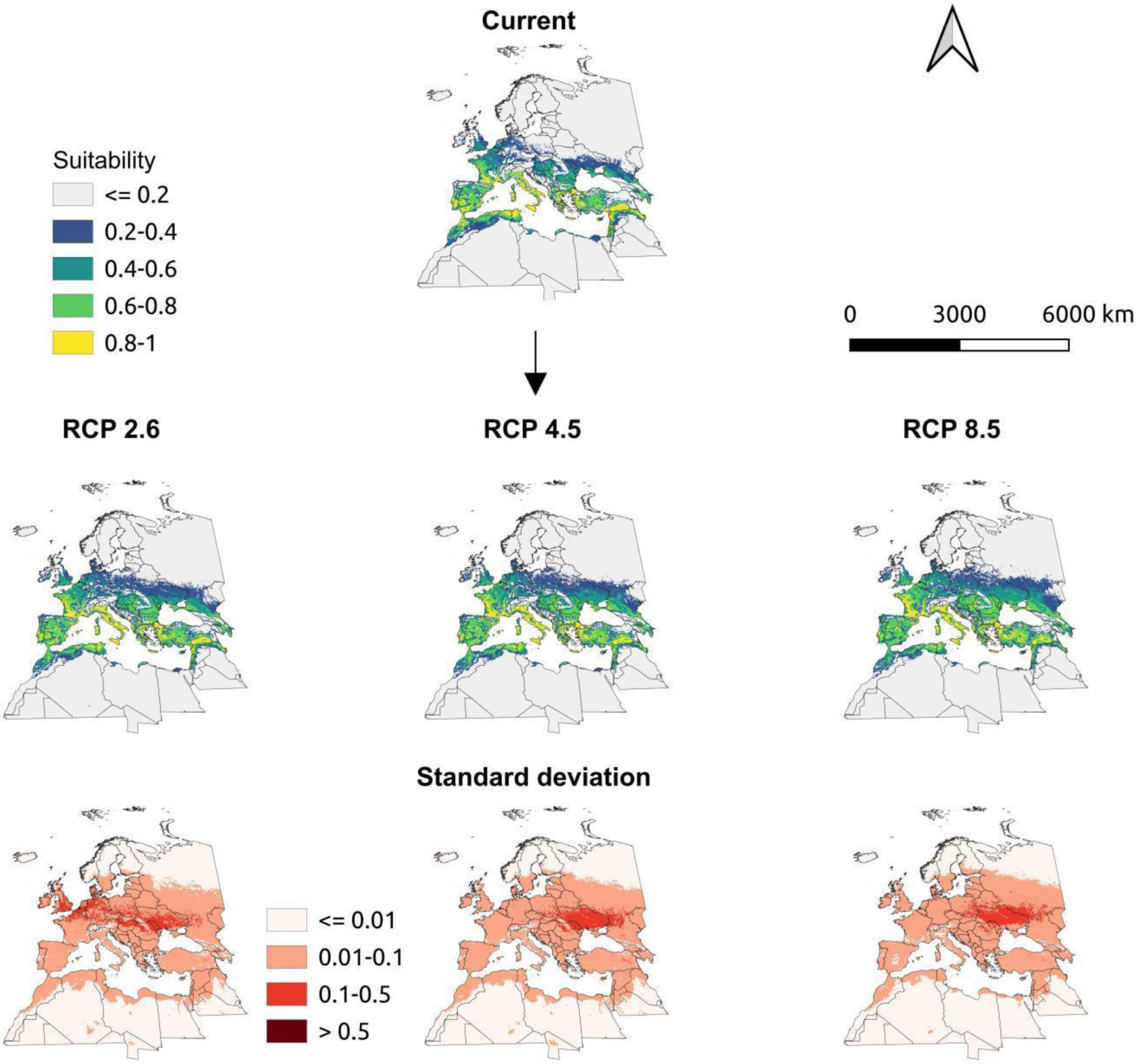
Current and future habitat suitability for the Eurasian Stone-curlew (*Burhinus oedicnemus*). Maps of current and future (2041-2060) breeding habitat suitability for the Eurasian Stone-curlew (*Burhinus oedicnemus*) in the Western Palearctic, under three Representative Concentration Pathways (RCP 2.6, RCP 4.5, RCP 8.5). The mean predicted suitability between five General Circulation Models (GCMs: CESM1-CAM5, FIO-ESM, IPSL-CM5A-MR, MIROC5, MPI-ESM-MR) is shown for each RCP, together with standard deviation between predictions according to the different GCMs. The projection used is World Mollweide (ESRI: 54009).

**Table 1.** Evaluation of Maxent species distribution models for the Eurasian Stone-curlew (*Burhinus oedicnemus*) in the Western Palearctic. Different values of beta (regularization multiplier) and different combinations of features were tested to tune model hyperparameters. The following metrics from a four-fold spatial block cross validation are reported: Area Under the ROC (Receiver Operating Characteristic) Curve of the test dataset (AUC_test_), the difference between AUC on the training dataset and AUC_test_ (AUC_diff_); Continuous Boyce Index (CBI). Standard deviations are reported in the adjacent columns. Results are ordered based on increasing overfitting (AUC_diff_). The model representing the best compromise among evaluation metrics is shown in bold.

| ID | Features | Beta | $AUC_{test}$ | SD | $AUC_{diff}$ | SD | CBI | SD |
| --- | --- | --- | --- | --- | --- | --- | --- | --- |
| **1** | **LQ** | **1** | **0.872** | **0.081** | **0.057** | **0.088** | **0.990** | **0.008** |
| 2 | LQ | 3 | 0.861 | 0.087 | 0.060 | 0.093 | 0.977 | 0.022 |
| 3 | LQH | 4 | 0.874 | 0.090 | 0.060 | 0.097 | 0.989 | 0.008 |
| 4 | LQH | 2 | 0.877 | 0.089 | 0.061 | 0.095 | 0.994 | 0.004 |
| 5 | LQH | 5 | 0.872 | 0.091 | 0.062 | 0.098 | 0.991 | 0.006 |
| 6 | LQH | 6 | 0.869 | 0.091 | 0.063 | 0.098 | 0.993 | 0.005 |
| 7 | LQH | 6 | 0.869 | 0.091 | 0.063 | 0.098 | 0.993 | 0.005 |
| 8 | LQH | 7 | 0.867 | 0.091 | 0.063 | 0.098 | 0.992 | 0.006 |
| 9 | LQH | 8 | 0.865 | 0.089 | 0.064 | 0.096 | 0.993 | 0.003 |
| 10 | LQH | 8 | 0.865 | 0.089 | 0.064 | 0.096 | 0.993 | 0.003 |
| 11 | LQ | 5 | 0.840 | 0.091 | 0.074 | 0.091 | 0.971 | 0.020 |
| 12 | LQ | 5 | 0.840 | 0.091 | 0.074 | 0.091 | 0.971 | 0.020 |
| 13 | LQ | 6 | 0.837 | 0.091 | 0.076 | 0.088 | 0.969 | 0.018 |
| 14 | LQ | 7 | 0.834 | 0.090 | 0.077 | 0.085 | 0.971 | 0.015 |
| 15 | LQ | 7 | 0.834 | 0.090 | 0.077 | 0.085 | 0.971 | 0.015 |
| 16 | LQ | 8 | 0.832 | 0.089 | 0.077 | 0.083 | 0.966 | 0.015 |

**Table 2.**
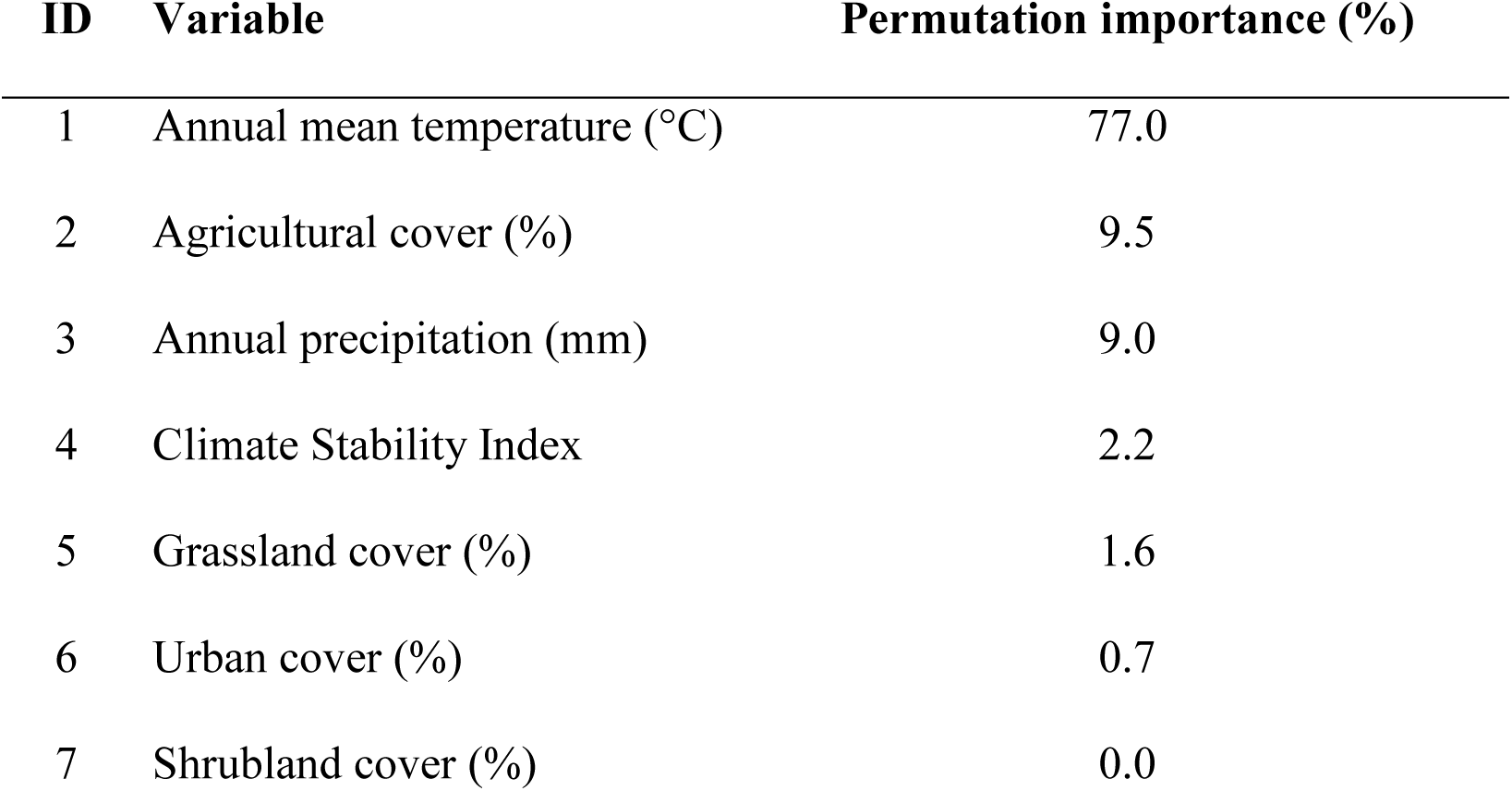
Environmental variables included in the tuned model used to predict current and future breeding habitat suitability for the Eurasian Stone-curlew (*Burhinus oedicnemus*) in the Western Palearctic, and their permutation importance. Permutation importance for a given variable is equivalent to the drop in AUC (Area Under the ROC - Receiver Operating Characteristic - Curve) after removing the variable and is expressed as percentage.

| ID | Variable | Permutation importance (%) |
| --- | --- | --- |
| 1 | Annual mean temperature (°C) | 77.0 |
| 2 | Agricultural cover (%) | 9.5 |
| 3 | Annual precipitation (mm) | 9.0 |
| 4 | Climate Stability Index | 2.2 |
| 5 | Grassland cover (%) | 1.6 |
| 6 | Urban cover (%) | 0.7 |
| 7 | Shrubland cover (%) | 0.0 |

### 3.2 Gap analysis

The 26.48 % of in-situ climate refugia for the Stone-curlew fell within the World Database on Protected Areas. In particular, such areas mostly overlapped with the currently suitable breeding areas (Figure 4: see also Figure 3 for the currently suitable areas). Large patches of in-situ climate refugia with a scarce cover of protected areas were mainly located in Northern Africa (Morocco, Algeria, Tunisia) and the Middle East (Israel, Turkey; Figure 4).

**Figure 4.**
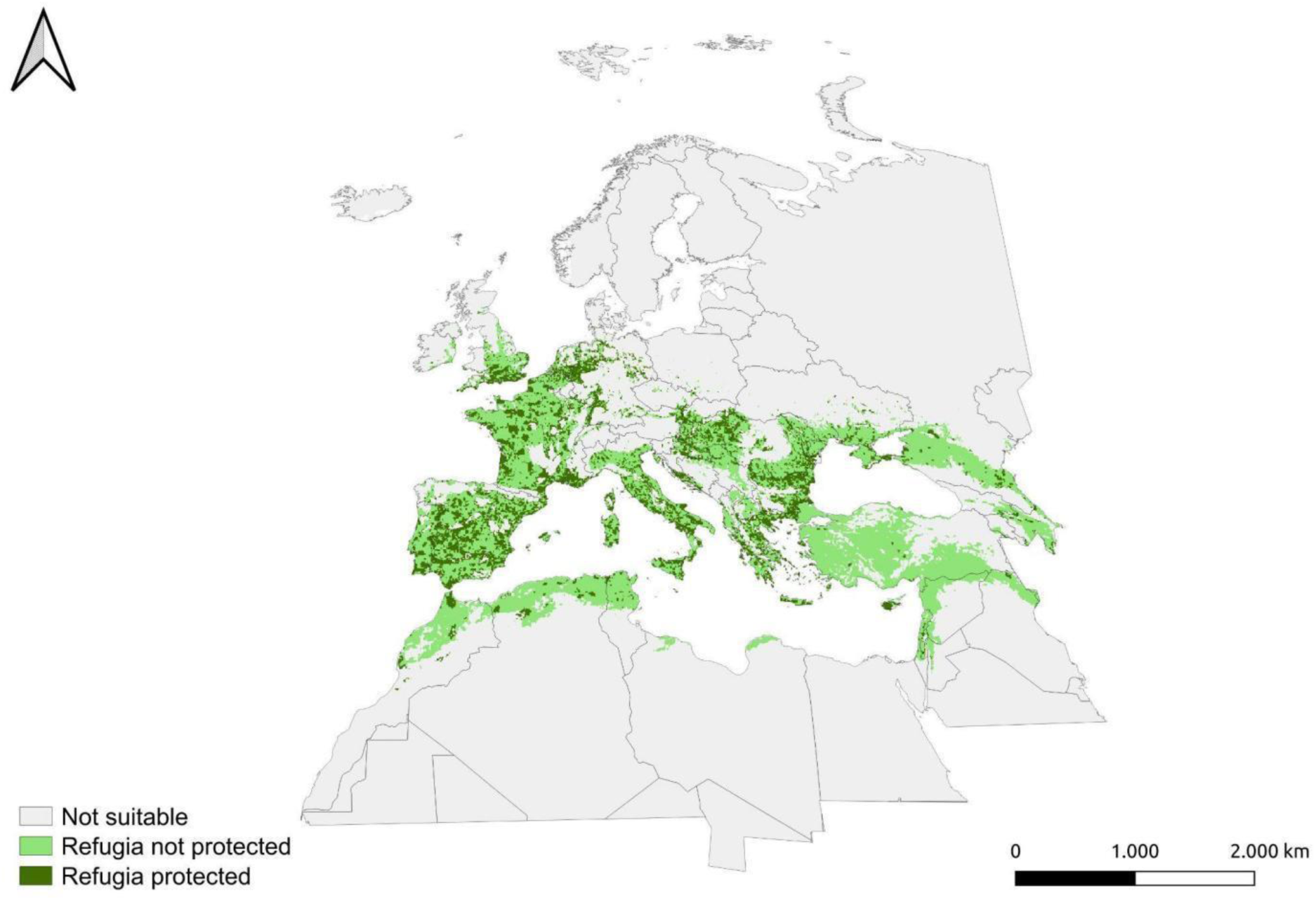
Coverage of in-situ climate refugia for the Eurasian Stone-curlew (*Burhinus oedicnemus*) by protected areas. In-situ climate refugia are areas predicted as suitable under both current and all future scenarios by a tuned Maxent species distribution model. Climate refugia inside the World Database on Protected Areas (www.protectedplanet.net) are shown in dark green, those outside in light green. The projection used is World Mollweide (ESRI: 54009).

## 4. DISCUSSION

Relying on spatially independent Stone-curlew’s occurrences, we developed a robust predictive framework for the species under climate change, based on the explicit tuning of Maxent models. The tuned model predicted an expansion of suitable breeding areas for the species towards the North-East under all future scenarios and allowed the identification of in-situ climate refugia, which fall mostly outside the current system of protected areas.

### 4.1. Environmental niche

Response curves showed results consistent with the knowledge on the autecology of species. The preference of the species for mild/warm climates is known from literature (Keller et al., 2020). The higher suitability at mid/high precipitation levels might be linked to the numerous breeding areas found in Mediterranean areas, that are rather arid sites despite relatively high annual precipitation due to a strong seasonality, with precipitations occurring primarily in autumn (Deitch et al., 2017). The very little permutation importance of urban cover suggests that urban areas are used mainly for occasional foraging and breeding by Stone-curlews, despite more frequent sightings in peri-urban settings, such as the area of Fiumicino Airport in Rome (Biondi et al., 2015). The choice of breeding sites primarily depends on the presence of extensive agricultural or natural areas nearby (forse è opportuna una citazione). Notably, the use of agricultural areas for breeding is increasingly documented (Issa and Muller, 2015), and agricultural areas might be good surrogates of natural open habitats (Figure 2). Stone-curlews have been reported to forage in agricultural areas with very different characteristics, but they require specific conditions for nesting that are not provided by areas with intensive agricultural practices (MacDonald et al., 2012). This calls for the adoption of agricultural practices compatible with Stone-curlew conservation needs, especially in areas close to where the species breeds and that are predicted to become suitable in the future. Furthermore, a Common Agricultural Policy that benefits specialised taxa instead of generalist one (Assandri et al., 2019) and that does not support intensive farming would benefit the conservation of the Stone-curlew.

### 4.2. Future breeding habitat suitability

The northward shift of suitable areas predicted by our model is consistent with previous projection for the species (Huntley et al., 2007; Keller et al., 2020), and with results obtained for other pseudo-steppic species, such as Great *Otis tarda* and Little Bustard *Tetrax tetrax* (Estrada et al., 2016) and European Roller *Coracias garrulus* (Kiss et al., 2020). The loss of suitable areas in the southern Mediterranean region (e.g., Northern Africa) is not surprising as the Mediterranean biogeographical region represents a hotspot of global change, and extensive biodiversity losses have been predicted here (Sala et al., 2000). Dispersal ability, biotic interactions and the carrying capacity of suitable habitats might determine whether a stable/increasing mean breeding habitat suitability translates into a stable/increasing population size (Bateman et al., 2013; Holloway et al., 2016). Habitat suitability has been linked to population size and trends in birds (Green et al., 2008; Stiels et al., 2021). We showed that large areas of the Western Palearctic are predicted to remain or become suitable in the future. Hence, the species may be able to maintain viable populations in the region. However, models predicted decreases of breeding suitability for the species at the southern edge of its distribution, so that marginal populations might need to track their niche in the future. On the one hand, evidence of the ability of animals to track their niche is contrasting (Chen et al., 2011; Devictor et al., 2008) and many terrestrial organisms have been shown to shift their distribution at a sufficient pace to track recent temperature changes (Chen et al., 2011). On the other side, species can also respond to global change persisting under unfavourable conditions being phenotypically plastic and becoming locally adapted (Valladares et al., 2014). An important aspect to consider is the effect of the length of night on the ability to successfully forage, as the Stone-curlew forages primarily during night (Caccamo et al., 2011) and the shorter length of nights at northern latitudes during breeding seasons can limit the ability to forage. A successful northward expansion would therefore imply a plastic shift towards foraging in daily hours or increased productivity to offset the limited foraging time. Finally, biological interactions and movement/dispersal constraints might prevent Stone-curlews from colonizing newly suitable areas (Bateman et al., 2013; Brambilla et al., 2020; Holloway et al., 2016). However, the observed variability in large-scale movements both within and between Mediterranean populations (Falchi et al., 2023) might suggest a significant potential of the species for colonizing new suitable areas.

### 4.3. Gap analysis

The Stone-curlew is philopatric in the breeding areas (Green, 1990) and this might delay habitat-tracking under environmental change. Therefore, in-situ climate refugia might act as a stronghold for the species, and anticipatory conservation efforts should primarily focus on these areas (Thuiller et al., 2019). Alongside, ensuring high connectivity conditions for the species in areas predicted to become unsuitable might contribute to maintain viable populations and facilitate niche tracking (Heller and Zavaleta, 2009). This is especially true considering that habitat fragmentation contributed to the species’ decline in the ‘90s (Tucker and Evans, 1997). Finally, enhanced monitoring efforts in areas predicted to become suitable might increase early-detection probability and allow to define and implement strategies to mitigate the negative effects of biotic interactions. For instance, our model evidenced the importance of agricultural areas, a LULC class heavily affected by farming and harvesting activities. Many in-situ climate refugia are located in intensively cultivated areas, corresponding to the ‘core of EU continental agriculture’ (D’Amico et al., 2013), namely England and the Po Plain, and extensively influenced by the Common Agricultural Policy reforms (Assandri et al., 2019). Furthermore, in France, over 60% of breeding pairs are found in the Central/Western region within arable crops (Issa and Muller, 2015; Malvaud and Blanchon, 1996). In these ecosystems, *ad hoc* management interventions on a local scale can be effectively used to favour Stone-curlew’s presence (Hawkes et al., 2021). A rather low percentage of the in-situ climate refugia are located within the framework of protected areas. Despite the predicted increase of suitable areas, due to the heavy reliance of the Stone-curlew on ecosystems affected by humans, it is critical to implement conservation measures and secure natural strongholds of the species towards colder climates (e.g., in northern France or Great Britain). Besides, pseudo-steppic species at large may benefit from protected areas (Santana et al., 2014) and thus an increased protection of in-situ climate refugia for the Stone-curlew, especially in areas with large gaps (North Africa, Middle East), might have a positive impact on other species.

### 4.4. Study limitations and future developments

The use of correlative species distribution models may present some limitations. First, SDMs assume an equilibrium condition between the species and the environment (Araújo and Pearson, 2005). Areas that have been abandoned by the Stone-curlew during the last decades of the past century may be recolonized, as happened in the UK following the targeted conservation efforts of the LIFE11INF/UK000418 ‘Securing the future of the Stone-curlew throughout its range in the UK’. This suggests that suitable climate and LULC conditions at the relatively coarse scale of our study exist in those areas. Thanks to constant improvements in climate and LULC scenarios, in the future, our workflow might be applied at a finer temporal scale for the Stone-curlew and, more generally, for steppic species and might include scenarios of LULC besides standard climatic change scenarios. Finally, we limited our study to the breeding period: understanding the drivers and changes of habitat suitability for the Stone-curlew across migratory and wintering grounds might provide further insights for its conservation.

## 5. CONCLUSIONS

This study refines current knowledge on the effects of climate change for the Stone-curlew, providing robust predictions of breeding habitat suitability for the species at an ecologically relevant spatial resolution, and conservation planners might benefit from our results by incorporating indications on the most relevant conservation areas in the development of action plans for the species. Our results might as well apply to steppe-land birds in general, and we advocate the use of a similar framework to model future habitat suitability for other sensitive species.

## ACKNOWLEDGEMENTS

We thank Diego Rubolini and Luca Forneris for technical help, Danae Portolou and Nikos Tsiopelas (Hellenic Ornithological Society) for providing aggregated occurrence data from Greece. D. Giunchi thanks all the students and collaborators who helped him in the field. We also thank Saverio Gatto for granting us the permission to use his Stone-curlew’s photograph in Figure 1 of the paper.

## SUPPLEMENTARY MATERIALS, DATA AND CODE

The supplementary materials, data and variables used to perform analyses, together with the R code to replicate them and the knitted document generated by R Markdown are found at: https://doi.org/10.5281/zenodo.11097919. Please note that occurrences from the UK provided by the British Trust for Ornithology are not included in the data as requested by the signed agreement of data concession.

## AUTHOR CONTRIBUTIONS

AS: conceptualization, data curation, formal analysis, investigation, methodology, supervision, writing - original draft, writing - review & editing, SR: conceptualization, data curation, formal analysis, investigation, methodology, writing - original draft, writing - review & editing, MF: formal analysis, methodology, writing - review & editing, MB: formal analysis, methodology, writing - review & editing, AMar: data curation, writing - review & editing, AMas: conceptualization, formal analysis, investigation, methodology, supervision, writing - review & editing, DG: conceptualization, data curation, formal analysis, investigation, methodology, supervision, writing - review & editing.

## Declaration of interests

☒The authors declare that they have no known competing financial interests or personal relationships that could have appeared to influence the work reported in this paper.

☐The authors declare the following financial interests/personal relationships which may be considered as potential competing interests:

## Notes

https://doi.org/10.5281/zenodo.11097919

### Competing Interest Statement

The authors have declared no competing interest.

